# Identification and full-genome assembly of novel Colorado potato beetle iflavirus and solinvi-like virus with SISPA Sequencing

**DOI:** 10.1101/2023.10.21.563391

**Authors:** Maria Antonets, Sergei Bodnev, Ulyana Rotskaya, Elena Kosman, Tatyana Tregubchak, Tatyana Bauer, Mamedyar Azaev, Vadim Kryukov, Denis Antonets

**Affiliations:** State Research Center of Virology and Biotechnology “Vector”, Rospotrebnadzor, 630559, Koltsovo, Russia; Novel Software Systems LLC, 630090, Akademika Lavrentiev ave. 6, Novosibirsk, Russia; Institute of Systematics and Ecology of Animals SB RAS, 630091, Frunze str. 11, Novosibirsk, Russia; Novosibirsk State University, 630090, Pirogova str. 2, Novosibirsk, Russia; MSU Institute for Artificial Intelligence, 119192, Lomonosov ave. 27, Moscow, Russia

## Abstract

The Colorado potato beetle is one of the most devastating potato pests widespread in the world. However, its viral pathogens remain highly unexplored. Here using SISPA high-throughput sequencing of Colorado potato beetle (CPB) samples derived from prepupal larvae that died from an unknown infection, we have identified two previously unknown viruses and assembled their full-length genomic sequences. The subsequent genetic and phylogenetic analysis of the obtained sequences demonstrated that the isolated viruses, named *Leptinotarsa iflavirus 1* and *Leptinotarsa solinvi-like virus 1*, are the novel representatives of *Iflaviridae* and *Solinviviridae* viral families, respectively. To the best of our knowledge, these are the first sequencing-confirmed insect viruses derived directly from CPB samples. And we also propose that *Leptinotarsa iflavirus 1* may be associated with lethal disease in CPB.

## Introduction

The Colorado potato beetle (CPB) – *Leptinotarsa decemlineata* of Chrysomelidae family – is one of the most widespread and destructive Solanaceae plants pests in the world. Although the use of insecticides helps to decline the beetle’s populations, the CPB is notorious for its ability to develop resistance to all chemicals registered to date [1]. That is why there is an urgent need for new effective methods to control CPB populations and also to develop new biocontrol strategies as the most environmentally friendly [2]. Currently, the existing methods of biological control of CPB populations use entomopathogenic fungi such as *Beauveria bassiana, Metarhizium sp*. [3] and bacteria *Bacillus thuringiensis* [4], but not viruses. However, entomopathogenic virus-based bioagents that are species-specific and generally safe for non-target organisms are valuable tools for developing integrated pest management strategies. As for other insect pests, positive examples include the nucleopolyhedroviruses of the *Baculoviridae* family, which are currently used to control a number of Lepidoptera species [4].

In a recent study we presented the results of metagenomic analysis of viral diversity in non-target CPB samples – namely, in public genomic and transcriptomic *Leptinotarsa decemlineata* NGS data obtained from the NCBI SRA database [5]. In these data, we have identified virus-associated genetic sequences belonging to more than 90 species and 32 families of viruses (excluding bacteriophages), a significant part of which were RNA viruses. It is worth noting that, targeted studies dedicated to the identification of CPB viruses have not been conducted to date.

When growing natural populations of the Colorado potato beetle under laboratory conditions [6,7], we observed larvae death rate of 8-35% (upto 60%) when they completed feeding and entered into the prepupal stage (7-11 days after molting at instar IV). The dead larvae were characterized by specific phenotype with body turgor loss, body straightening and legs stretching (Fig. 1). 1-3 days post mortem the hemolymph acquired a dark tint characteristic to septicemia (Fig. 1C). To the best of our knowledge these particular disease specific symptoms were not described earlier in CPB. In the samples of dead CPB prepupae we identified two novel RNA viruses, one of which was attributed to *Iflaviridae* family, and the second – to *Solinviviridae* family.

**Fig. 1.**
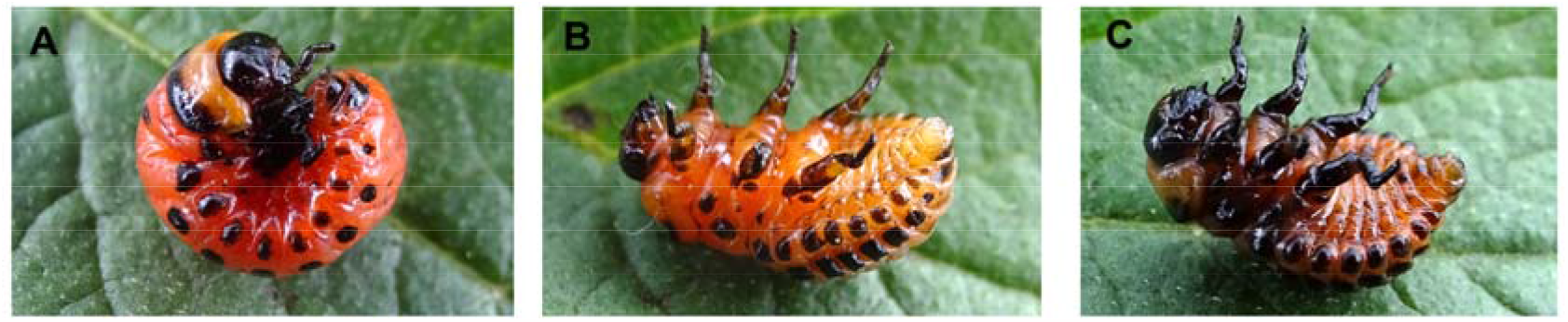
Disease symptoms. A — the healthy prepupa; B — the prepupa 1 day after death; C — the prepupa 3 days after death.

Iflaviruses belong to *Iflaviridae* family of order Picornavirales, and are characterized by a single-stranded RNA+ genome with a length of 9-11 thousand nucleotides. Their genome contains a single open reading frame encoding a polyprotein that is proteolytically processed into functional viral proteins [8]. More than 550 whole genome iflavirus sequences have been published to date to the NCBI GenBank database, of which 134 were published in 2021, 105 in 2022 and 42 in 2023. Iflaviruses infect arthropods with most of the described representatives infect insects. Iflaviruses target specific tissues of the insect host, often affecting crucial physiological processes. They can oppose the host immune responses by interfering with antiviral defense mechanisms and ensuring successful viral replication [9]. In some cases iflaviruses cause persistent infection, which makes them particularly interesting [10].

Solinvi-like viruses belong to *Solinviviridae* family of order Picornavirales. The *Solinviviridae* family consists of positive-sense RNA viruses with non-segmented genomes that are approximately 10-11 kb in length. Interestingly, their capsid proteins are encoded towards the 3′-end of the genome, where they can be expressed both from a subgenomic RNA and as a replication polyprotein extension, which is unusual [11]. *Solinviviridae* viruses infect arthropods, with most described representatives infecting insects and crustaceans [11,12]. Some of these viruses are known to cause chronic infections and others, which are more virulent, cause systemic infections and acute mortality, however little is currently known about the effects of the most of these viruses on their hosts [12]. Only about 30 whole genome sequences of *Solinviviridae* viruses have been published to NCBI GenBank database to date.

## Results and Discussion

### Assessment of viral load with quantitative PCR

In our previous analysis of public CPB NGS data we were able to assemble several near full-length genomes of viruses related to *Iflaviridae* family [5] and thus we proposed that the observed CPB prepupal larvae death might be at least partially attributed to iflaviral infection. To test this possibility, we undertook a qPCR study to assess the quantities of viral genetic material in symptomatic dead prepupae in comparison to asymptomatic healthy ones. The viral RNA-dependent RNA polymerase (RdRp) gene was selected to assess the viral load and *L. decemlineata* ribosomal protein RP4 and RP18 genes were taken as reference ones, as it was proposed in [13]. The designed oligonucleotide primers are shown in Table 1. The qPCR study demonstrated the increased levels of viral RdRp in fresh dead symptomatic larvae (“infected”) as compared to healthy asymptomatic ones (see Fig. 1 A, B) – the mean relative RdRp expression in “infected” group was about 32.9 times higher than in control (Fig. 2). And the observed difference was found to be statistically significant (p = 0.0357).

**Table 1.**
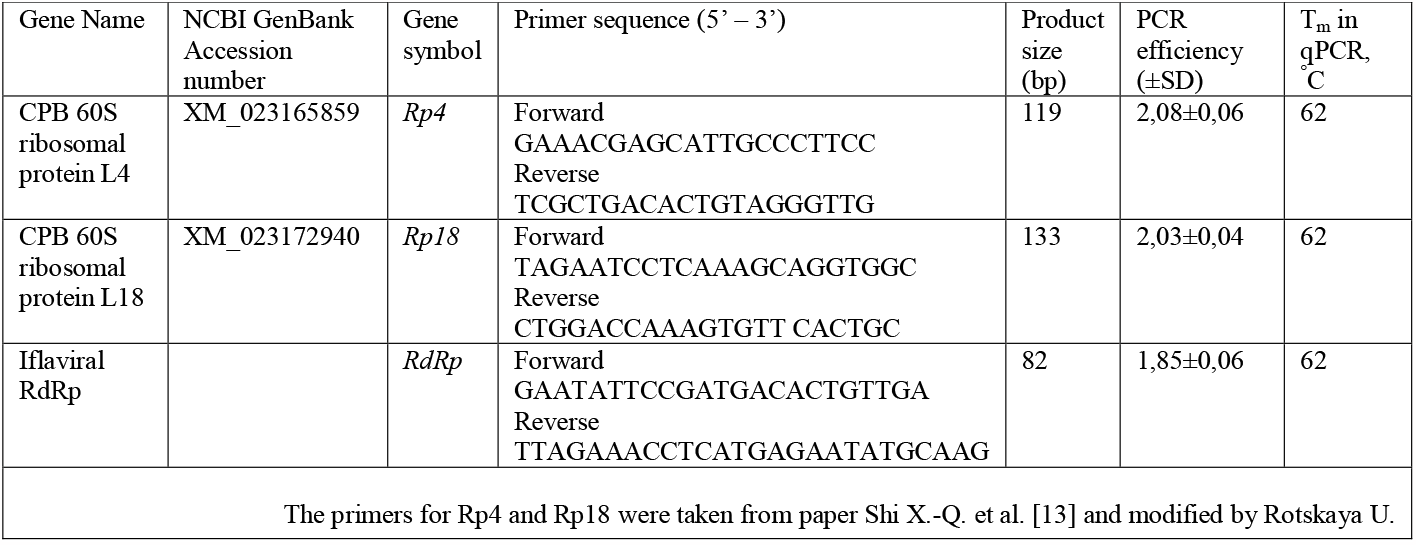
The selected genes and oligonucleotide primer sequences used for qPCR.

**Fig. 2.**
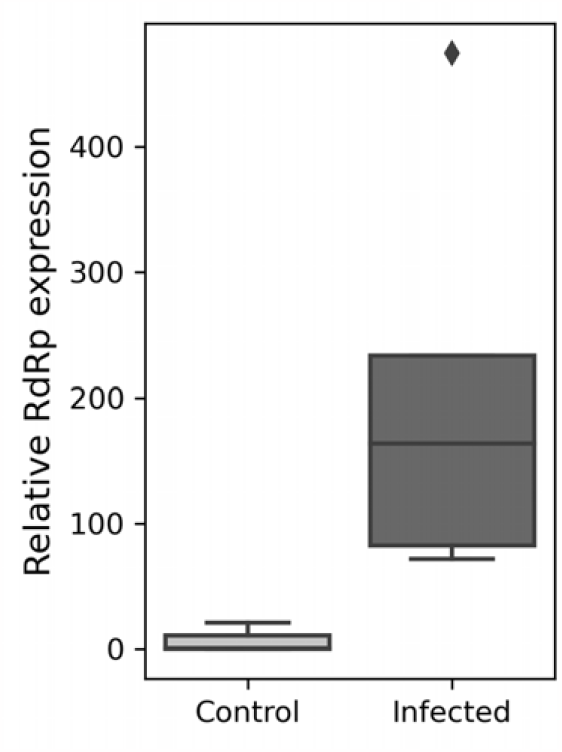
Iflaviral RdRp relative expression in infected and control samples of CPB prepupae. The relative RdRp expression was determined as detailed in corresponding Material and Methods section. The RdRp expression values were normalized to reference genes: ribosomal proteins L4 and L18 (Rp4 and Rp18). The difference in viral loads was found to be statistically significant (Mann-Whitney two-tailed test, p = 0.0357).

Thus, the significantly increased levels of viral RdRp were observed in tissues of freshly dead larvae with characteristic symptoms, which indicates active production of viral particles. Unfortunately, in current study it was impossible to attribute the larvae death to this particular virus infection. To prove that the virus causes lethal infection in CPB, it is necessary to purify the viral particles in order to determine their infectivity and lethal dose in further *in vitro* and *in vivo* studies. And it is extremely important to determine the complete genomic sequence of the target virus.

### Sequencing and assembly of viral genome sequences

The sequencing yielded 473.5×2 thousand reads, with an average length of 124 nt, for the CPB3 sample and 480.5×2 thousand reads, with an average length of 127 nt, for the CPB6 sample. The *de novo* assembly yielded a full genome sequence of an iflavirus named *Leptinotarsa iflavirus 1* (in sample CPB3) with an average coverage depth of 834x (7.1 % reads) and a full genome sequence of solinvi-like virus named *Leptinotarsa solinvi-like virus 1* (in sample CPB6) with an average coverage depth of 1116x (10.87 % of reads). The genetic sequences of *Leptinotarsa iflavirus 1* and *Leptinotarsa solinvi-like virus 1* were deposited to GenBank (with accession IDs *OR613011* and *OR613010*, respectively).

### Analysis of viral genetic sequences with InterProScan

The *Leptinotarsa iflavirus 1* genome sequence is 9887 nt long, excluding the polyA-tail, and contains a single ORF (positions 561-9707), 5’-UTR (560 nt) and 3’-UTR (180 nt). The ORF encodes a polyprotein of 3049 aa, which, according to the results of InterProScan, contains domains characteristic to iflaviruses and their mutual arrangement is also typical for the *Iflaviridae* family. These domains are the following: two rhv-like domains in two subunits of the Picornavirus/Calicivirus-like capsid protein (*IPR033703*, E-values of 5.96×10^-13^ and 1.15×10^-21^, respectively), domain belonging to Calicivirus-like capsid proteins superfamily (*IPR029053*, 2.5×10^-32^), RNA helicase domain (*IPR000605*, 1.4×10^-18^), domain belonging to cysteine 3C peptidases superfamily (*IPR009003*, 2.36×10^-16^) and RNA-dependent RNA polymerase domain (*IPR043502*, 4.65×10^-78^) (Fig. 3A). The amino acid sequence of the *Leptinotarsa iflavirus 1* polyprotein has the highest homology with *Apis flavirus 2* polyprotein (*UCR92484*); according to the BLASTp results their mutual identity is 37.72 %.

**Fig. 3.**
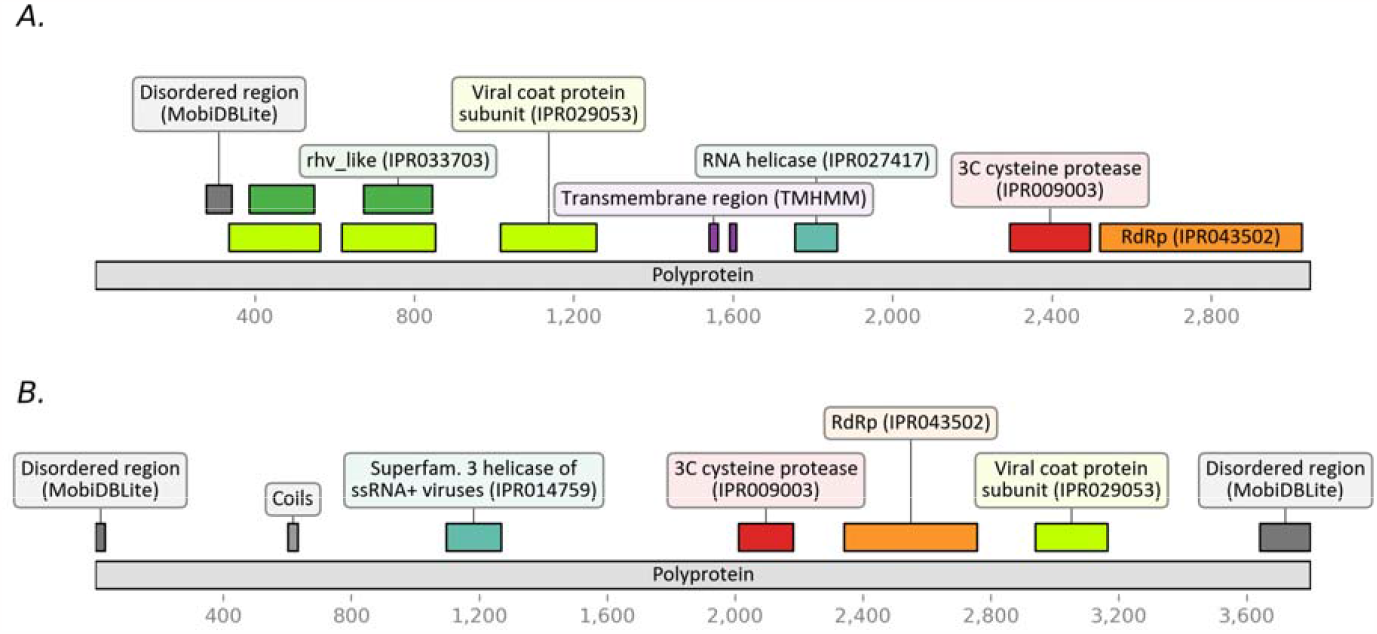
The results of viral polyproteins annotation with InterProScan. (A) the polyprotein of *Leptinotarsa iflavirus 1* (*WNV56445*) and (B) *Leptinotarsa solinvi-like virus 1* polyprotein (*WNV56444*).

The genome sequence of *Leptinotarsa solinvi-like virus 1* is 11603 nt long, excluding the polyA tail, and contains a single ORF (coordinates 62-11458). This ORF encodes a polyprotein of 3800 aa, which, according to InterProScan, contains the following domains: RNA helicase domain (*IPR000605*, E-value: 7.2×10^-20^), a domain belonging to the superfamily of cysteine 3C peptidases (*IPR009003*, 7.41×10^-6^), RNA-dependent RNA polymerase domain (*IPR043502*, 9.39×10^-55^) and Calicivirus-like capsid protein domain (*IPR029053*, 1.5×10^-18^) (Fig. 3B). The amino acid sequence of *Leptinotarsa solinvi-like virus 1* polyprotein has the highest homology with the polyprotein of *Hangzhou Solinvi-like virus 1* (*UHR49784*), derived from samples of *Altica cyanea* leaf-feeding beetle (Chrysomelidae). According to the BLASTp results, their mutual sequence identity is 33.32 %.

### Phylogenetic analysis of viral RdRp proteins

Since iflaviral sequences are very diverse, the phylogenetic analysis was performed using the most conserved polyprotein fragments containing RdRp. The multiple amino acid sequence alignment was built using the sequences with the highest homology to *Leptinotarsa iflavirus 1* RdRp (at least 30% identity) selected from NCBI nr database using BLASTp. The phylogenetic tree was constructed with IQ-Tree software using the maximum likelihood method (Fig. 4). The sequences closest to *Leptinotarsa iflavirus 1* RdRp were the RdRp of *Apis iflavirus 2* (*UCR92484*), derived from metagenomic samples of honey bees collected in China in the Henan province in 2017, and the RdRp of *Lampyris noctiluca iflavirus 1* (*QBP37019*), obtained from metagenomic analysis of a firefly sample collected in Finland in 2017.

**Fig. 4.**
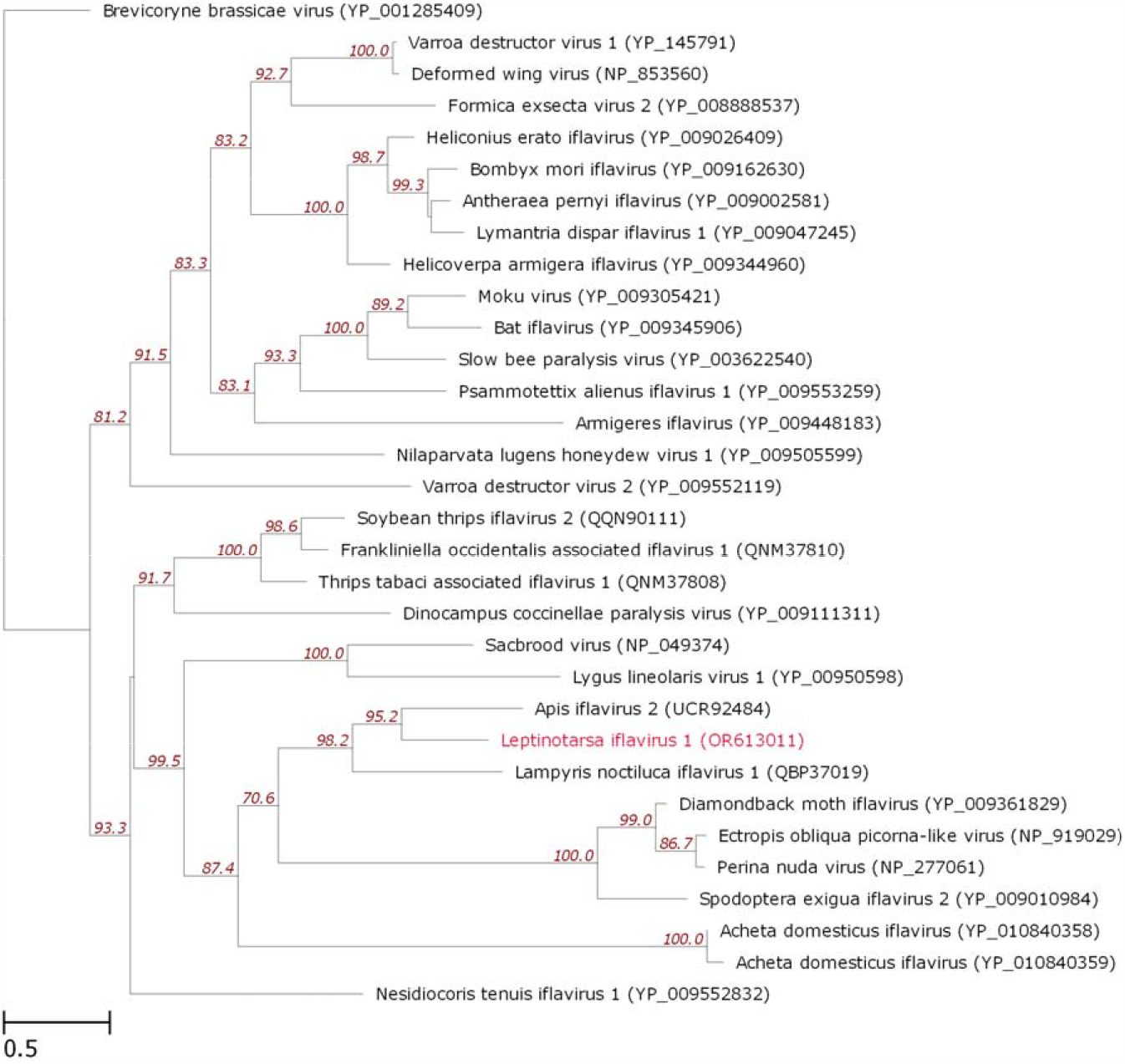
The phylogenetic tree of RdRp amino acid sequences of *Leptinotarsa iflavirus 1* and closely related iflaviruses and picorna-like viruses. The tree was constructed with IQ-Tree by the maximum likelihood method. The numbers next to the branching points denote the support indices (>= 70 %) determined by boot strap statistical analysis (1000 replications). The tree is drawn to scale, with branch lengths corresponding to evolutionary distances.

To construct the phylogenetic tree of *Leptinotarsa solinvi-like virus 1* RdRp, we used the multiple RdRp amino acid sequence alignment taken from ICTV (https://ictv.global/sites/default/files/inline-images/Figure3-1.v2.aa.alignment.fst, accessed on August 2023) [11]. The alignment was augmented with sequences closely related to *Leptinotarsa solinvi-like virus 1* RdRp extracted from NCBI nr database with BLASTp (with at least 30 % identity). Fig. 5 demonstrates a fragment of the resulting phylogram produced with IQ-Tree.

**Fig. 5.**
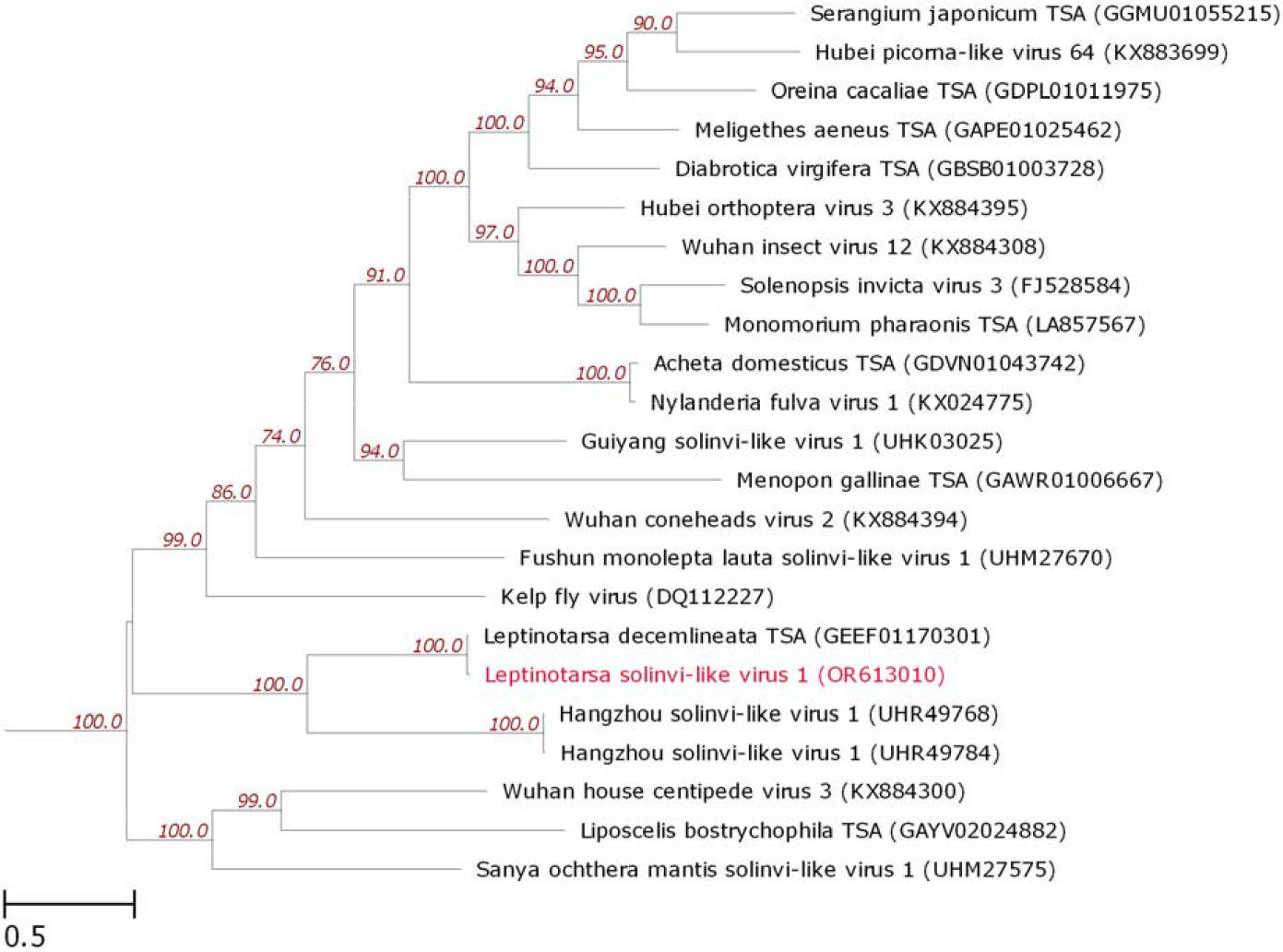
The phylogenetic tree of RdRp amino acid sequences of *Leptinotarsa solinvi-like virus 1* and closely related solinviviruses and solinvi-like viruses. The tree was constructed with IQ-Tree by the maximum likelihood method. The numbers next to the branching points denote the support indices (>= 70 %) determined by boot strap statistical analysis (1000 replications). The tree is drawn to scale, with branch lengths corresponding to evolutionary distances. The tree is a part of a bigger one (not shown).

The *Solinviviridae* sequences extracted from ICTV were also found to include RdRp encoded by the putative viral genetic sequence Leptinotarsa decemlineata TSA (GenBank: *GEEF01170301*), obtained from transcriptomic assembly of CPB samples (BioProject ID: *PRJNA297027*). This RdRp has the highest homology with the *Leptinotarsa solinvi-like virus 1* RdRp among all the compared sequences; the nucleotide sequence identity was 97 % (1212/1251). We have also performed pairwise comparison of the full-length polyproteins encoded by *Leptinotarsa solinvi-like virus 1* genome and Leptinotarsa decemlineata TSA sequence. The total length of the polyprotein sequence encoded by Leptinotarsa decemlineata TSA is 3819 aa. The identity of the amino acid sequences of the compared polyproteins is 96.86 % (3699 out of 3819 aa), most amino acid substitutions were found to be conservative (the sequence homology according to BLOSUM62 equals to 98.56 %). As compared to the *Leptinotarsa solinvi-like virus 1* polyprotein, a fragment of 19 aa is inserted. The comparison of nucleotide sequences demonstrated that the Leptinotarsa decemlineata TSA sequence lacks terminal fragments, including the polyA tail. The polyprotein encoding nucleotide sequence of Leptinotarsa decemlineata TSA contains 480 SNPs, including 379 transitions, 101 transversions, 360 synonymous and 120 missense substitutions (with 55 being conservative), and 1 deletion (57 nt). Thus, it appears that the Leptinotarsa decemlineata TSA sequence (*GEEF01170301*) corresponds to a near full-length genomic sequence of a related solinvi-like virus. In addition to Leptinotarsa decemlineata TSA, the other closest to *Leptinotarsa solinvi-like virus 1* RdRp were the two RdRp sequences of *Hangzhou solinvi-like virus 1* (NCBI Protein: *UHR49768* and *UHR49784*), derived from metagenomic samples of *Altica cyanea* leaf-feeding beetle (Chrysomelidae) collected in China in 2016.

Thus, for the first time, using the SISPA-Seq method, the full-length genome sequences of viruses belonging to the order Picornovirales were obtained from natural samples of CPB. The phylogenetic analysis allowed to attribute these viruses as representatives of the iflaviruses and solinvi-like viruses – *Leptinotarsa iflavirus 1* and *Leptinotarsa solinvi-like virus 1*, respectively. The analysis of amino acid sequences of polyproteins encoded by the genomes of these viruses confirmed the presence of characteristic viral proteins and demonstrated the correspondence of their domain structure to that of other described iflaviruses and solinvi-like viruses, respectively.

However, we cannot be sure that the observed *L. decemlineata* larvae death was caused by these particular viruses, although the external symptoms (delayed septicemia) and qPCR data (near 30-fold increase of viral load as compared to healthy insects), as well as the detection of these particular viruses in tissues of dead symptomatic larvae with SISPA-Seq, speak in support of this hypothesis. Noteworthy is the death of CPB at a certain period of their development (at transition to the prepupa stage), and we assume this might be associated with increased insects’ vulnerability to generalized viremia at the beginning of body restructuring period. Further studies will be aimed at isolating the viral particles and assaying their infectivity and pathogenicity in detail using both *in vitro* and *in vivo* models. We believe that our results may contribute to future development of novel Colorado potato beetle biological control methods.

## Acknowledgements

This work was supported by the Ministry of Science and Higher Education of the Russian Federation (agreement # 075-15-2019-1665). Insect collection, their laboratory maintenance, tissue sample preparation and qPCR were supported by the Russian Scientific Foundation (grant # 22-14-00309).

## Materials and Methods

### Biological samples collection

*L. decemlineata* III-IV instar larvae were collected from private potato fields free from any insecticides treatment (Karasuk, Novosibirsk region, Russian Federation; 53°43′N; 77°38′E). Due to these potato fields not being located in protected areas, there was no need for special permission to collect beetles. The landowners did not prevent access to the fields. Endangered or protected species were not used in this work. The insects were kept in laboratory conditions as described earlier [6,7]. The larvae were kept in ventilated plastic containers (300 ml; 10 larvae per container) at temperature of 24-25 °C and 16:8 h light/dark period daily fed with fresh *Solanum tuberosum* foliage. Insects that died in the prepupal stage with presumable viral infection symptoms (Fig. 1) were immediately frozen in liquid nitrogen and then stored at –80 °C until RNA extraction.

### Samples preparation and qPCR

The whole larvae were frozen in liquid nitrogen and stored at –18 °C. The samples were lyophilized at –53 °C and 400 mTorr for 24 hours. The lyophilized corpses were homogenized by micro pestles in liquid nitrogen and treated with Lira-reagent (BioLabMix Ltd., Novosibirsk, Russia). The subsequent total RNA extraction was performed according to Lira-reagent protocol. DNase treatment was carried out according to DNase I (RNase-free) kit (TransGen Biotech Co. Ltd., China) protocol. The reverse transcription of RNA to cDNA was performed with RevertAidTM M-MuLV Reverse Transcriptase (Fermentas, Vilnius, Lithuania) and 2.0 pMol of 9N primers.

qPCR was performed as described earlier [14,15]. The PCR reaction conditions were the following: 95 °C for 3 min, followed by 40 cycles of 94 °C for 15 s and 64 °C for 30 s, followed by the melting curve analysis (70-90 °C). The *L. decemlineata* ribosomal protein RP4 and RP18 genes were taken as reference, as it was proposed in [13]. The viral RNA-dependent RNA polymerase (RdRp) gene was selected to assess the viral load. Primers were designed with Primer-BLAST tool [16] and IDT OligoAnalyser 3.1 [17]. The designed oligonucleotide primers were synthesized by Biosset Ltd. (Novosibirsk, Russia). The primer sequences are provided in Table.1.

### qPCR gene expression calculations and statistical analysis

Gene expression was calculated using the 2ΔΔCq method with Bio-Rad CFX manager software (Bio-Rad, USA). The program Past 4.03 was used for analyses qPCR data. The statistical significance of observed differences in normalized viral RdRp expression values in dead larvae samples with presumable viral infection (N=5) and in asymptomatic larvae samples (N=3) was examined with two-sided Mann-Whitney test (the critical significance level was set to 0.05). The statistical analysis was performed with R statistical analysis environment (v.4.2.1) [18].

### Samples preparation and sequencing

Individual dead CPB prepupae were homogenized in 1 ml of standard PBS manually in Eppendorf 1.5 ml microtubes using sterile Axigen disposable pestles (PES-15-B-SI, Axigen, USA). The homogenates were centrifuged on Eppendorf Minispin centrifuge at 10000g for 10 minutes, the supernatants were collected into 15 ml tubes and diluted 10 times with sterile PBS. The resulting solutions were filtered through a Millex-HV filter pad with a pore diameter of 0.45 nm (Millex® PVDF syringe filter, Merk, USA) and the filter pads were additionally washed with 5 ml of sterile PBS. The resulting solutions were concentrated to a volume of 200 μl using Amicon® Ultra 4 50K centrifuge filters (Merk Millipore Ltd, Ireland) in Eppendorf 5804 centrifuge at 3000g with adapters for 15 ml tubes. The aliquots of 100 μl of the resulting solutions were supplied with 2 μl of MgCl_2_ (100 mM), shaken and 10 μl of benzonase (Sigma, USA) with an activity of 250 units/μl was added. The suspension was incubated at 37 °C for 30 minutes and then EDTA was added to inactivate the benzonase.

Then total RNA was extracted from 100 μl of benzonase-treated sample with TriZol (ThermoFisher Scientific, USA) according to the recommended protocol. 40 μg of glycogen in the form of an aqueous solution with a concentration of 20 mg/ml was used as a co-precipitant. The precipitate was dissolved in 30 μl of deionized water, then the solution was used to obtain the double-stranded DNA fragments with SISPA (Sequence-Independent, Single-Primer Amplification) protocol, according to [19].

The resulting dsDNA fragments were purified from unspent components and reaction products with AMPure beads (Beckman Coulter), and then used to prepare NGS libraries. Nucleic acid concentrations were measured with Qubit 3.0 using the Qubit dsDNA HS Assay Kit (Thermo Fisher Scientific). The NGS library preparation was carried out using the NEBNext Ultra II FS DNA Library Prep Kit for Illumina (NEB, USA), which performs fragmentation, end repair and dA-tailing, and adapter ligation with a single enzyme mix. The NEBNext Multiplex Oligos for Illumina (Index Primer Set 1) (NEB, USA) was used for multiplexing. The sequencing was performed on Illumina iSeq 100 platform (300 cycles).

### Sequencing Data Availability

The resulting FASTQ files were deposited to SRA with corresponding IDs: *SRR25929780, SRR25929781, SRR25929782, SRR25929783, SRR25929784* (BioProject: *PRJNA1013476*). The data was made public on October 11, 2023.

### Bioinformatic Analysis

The FastQ files obtained from sequencing were processed with fastp v0.20.0 [20] to remove adapters, and short and low-quality sequences (with quality less than 20 and with length of no more than 30 nt). The preprocessed reads were assembled into contigs using MEGAHIT v1.2.9 assembler [21] with the meta-sensitive parameter enabled. The resulting contigs were analyzed using BLASTx v2.9.0+ [22] against the NCBI nr database (https://ftp.ncbi.nlm.nih.gov/blast/db/FASTA/nr.gz, accessed on August 2023). The quality of assembled viral contigs was checked by back-aligning the obtained reads with BWA MEM v0.7.17-r1188 [23]. Target viral sequences were translated in 6 translation frames with EMBOSS transeq v6.6.0.0 [24]. The open reading frames within the polyproteins were determined by homology with BLASTp v2.9.0+ (against NCBI nr database). The polyproteins domain annotation was carried out with InterProScan [25]. The genomic sequences annotations were visualized using the DnaFeaturesViewer v3.1.2 Python3 package [26]. The phylogenetic analysis of *Leptinotarsa iflavirus 1* and *Leptinotarsa solinvi-like virus 1* RdRp proteins was performed using viral RdRp protein sequences with amino acid identity ≥ 30% selected with BLASTp from the NCBI nr database, and also the sequences taken from ICTV-provided alignment of *Solinviviridae* RdRp proteins (https://ictv.global/sites/default/files/inline-images/Figure3-1.v2.aa.alignment.fst, accessed on August 2023). The sequences were aligned with mafft v7.313 [27] using the L-INS-i algorithm with 1000 iterations. The phylogenetic trees were constructed using the maximum likelihood method with IQ-Tree v2.0.3 [28]. The following substitution models were used: LG+I+G4 for *Leptinotarsa iflavirus 1* and LG+F+R5 for *Leptinotarsa solinvi-like virus 1*. The selection of replacement models was carried out with ModelFinder algorithm [29]. Bootstrap analysis was performed with UFBoot algorithm [30] in 1000 replications. The phylogenetic trees were visualized with ete3 v3.1.2 Python3 package [31].

